# The viral vectored vaccine targeting Thymosin β10 controls tumour growth in a murine model of prostate cancer

**DOI:** 10.1101/2021.04.06.438563

**Authors:** Federica Cappuccini, Julius Mueller, Emily Pollock, Adrian VS Hill, Nicola Ternette, Irina Redchenko

## Abstract

Direct analysis of MHC-presented antigens by mass spectrometry has been successfully applied for the evaluation of the antigenic landscape of different human tissues and malignancies in the past. In the current study, for the first time this approach has been applied to investigate a murine prostate cancer peptidome for the development of next generation prostate cancer vaccines using the TRansgenic Adenocarcinoma of Mouse Prostate, TRAMP, model. We have performed immunopeptidomic analysis of the normal mouse prostate and mouse prostate adenocarcinoma samples and identified a number of peptides overexpressed on tumour tissue compared to the normal prostate. We selected two of these proteins previously reported to play a role in carcinogenesis, thymosin beta 4 (Thyβ4) and thymosin beta 10 (Thyβ10), expressed them in the chimpanzee adenovirus (ChAdOx1) and Modified Vaccinia Ankara (MVA) recombinant viral vectors and evaluated these constructs as a candidate prostate cancer vaccine in a murine prostate cancer model. We have found that the vaccine encoding the Thyβ10 polypeptide tethered to the MHC class II associated invariant chain, CD74, significantly delayed outgrowth of established tumours in an ectopic TRAMP-C1 mouse model of prostate cancer. The observed tumour-protective efficacy was not associated with Thyβ10-specific cellular or antibody responses, suggesting an existence of other mechanisms underlying anti-tumour effect in this model that are yet to be elucidated.

## 1. Introduction

Prostate cancer is the tumour type that is likely to be amenable to antigen-specific immunotherapy and the one for which the first therapeutic cancer vaccine, Sipuleucel-T, was approved by the US FDA in 2010 [1]. Despite a concerted effort of the cancer scientific community over the last decades, experimental vaccines against cancer, and prostate cancer in particular, have been modestly effective in terms of immunogenicity and clinical efficacy. There is a number of factors potentially underlying the vaccine relative inefficacy including tumour suppressive microenvironment, active adaptive mechanisms mounted by tumours in the face of rising immune response, and immune tolerance to tumour-associated self-antigens. In view of that, the choice of a target antigen(s) is one of the crucial decisions for designing a successful cancer vaccine. Sipuleucel-T and other experimental clinically tested prostate cancer vaccines target commonly shared prostate tumour specific antigens, e.g PAP, PSA, PSMA, PSCA, STEAP1 [2–5]. Their specificity for prostate tissue and, therefore, suitability as vaccine targets have been confirmed by immunohistochemistry (IHC) [6, 7] or gene expression analysis [8, 9]. However, IHC data do not provide information on the antigen processing and presentation and, therefore, the availability of molecular targets, namely the MHC-peptide complexes (MHCp), on the tumour cell surface for recognition by tumour-specific T cells. Mass spectrometry (MS) is the tool that can potentially overcome these limitations through the assessment of naturally processed and presented tumour antigens. In recent years, MS has become a driving force in the field of peptide identification and quantitation and is considered to be a valuable methodology to comprehensively interrogate the repertoire of MHCp presented naturally *in vivo* [10]. Immunopeptidomics analysis for tumour antigen discovery has been already performed for several human tumour types [11–17]. In the current study, for the first time this method has been applied to investigate naturally processed prostate tumour antigens for the development of next generation prostate cancer vaccines using a mouse model of prostate cancer, the transgenic adenocarcinoma of mouse prostate TRAMP [18]. We have performed immunopeptidomic analysis on the normal mouse prostate and mouse prostate adenocarcinoma samples and identified a number of peptides overexpressed on tumour tissue compared to the normal prostate. Some of the identified peptides were mapped to proteins previously reported to play a role in carcinogenesis. We selected two of these proteins containing epitopes presented on the surface of the TRAMP tumour but absent from a healthy murine prostate gland, thymosin beta 4 (Thyβ4) and thymosin beta 10 (Thyβ10), and expressed them in the chimpanzee adenovirus (ChAdOx1) and Modified Vaccinia Ankara (MVA) recombinant viral vectors for the subsequent evaluation of these constructs as a candidate prostate cancer vaccine in an ectopic TRAMP-C1 mouse model of prostate cancer.

## 2. Materials and Methods

### 2.1. Mice and Cell lines

Six-week-old male C57BL/6 mice used in this study were purchased from Harlan, UK, unless stated otherwise. Mouse care and experimental procedures were carried out in accordance with the terms of the UK Animals (Scientific Procedures) Act Project Licenses (PPL 30/2947, PPL P0D369534) and approved by the University of Oxford Animal Care and Ethical Review Committee.

TRAMP-C1 cell line was purchased from the American Type Culture Collection (ATCC) and maintained in high glucose DMEM supplemented with 5 % fetal bovine serum, 5 % NuSerum (Becton Dickinson), 4 mM L-Gln, 1 % Pen-Strep, 10 nmol/L dihydrotestosterone (Sigma), and 5 μg/mL bovine insulin (Sigma). Alternatively, TRAMP-C1 cells were cultured for 48 hours with recombinant mouse IFN-γ (100U/ml) and then gently scraped and rinsed in DPBS. After centrifugation, the cell pellet was stored at −20 °C until further lysis for mass spectrometry.

The hybridoma cell line HB-51 (ATCC) was cultured in a bioreactor and maintained in DMEM supplemented with 20 % ultra-low IgG fetal bovine serum, 4 mM L-Gln, 1% Pen-Strep. Cells were harvested and reseeded twice weekly; supernatant was collected and stored at −20 °C until further mAb purification.

### 2.2. Monoclonal antibody purification from hybridoma supernatant

Briefly, hybridoma supernatants were incubated for 1 hour at room temperature (RT) with Protein A resin. Resin was then collected in a column and washed with a minimum of 10 column volumes PBS. Monoclonal antibodies were eluted with 100 mM Glycine buffer, pH 2.5 directly into a reservoir of 1M Tris buffer, pH 9.2, for immediate neutralisation.

### 2.3. Generation of lysates from cells and tissue material

For analysis of mouse samples, one malignant TRAMP mouse prostate, pooled normal prostates from C57BL/6 wild type mice (total weight of pooled normal prostates equalled to the TRAMP malignant prostate), and 10^8^ IFN-γ treated TRAMP-C1 cells were processed in parallel. The cells and tissue material were homogenized in lysis buffer (1% Igepal, 300 mM sodium chloride, 100 mM Tris, pH 8.0) using a Dispomix homogenizer (Medic Tools AG, Miltenyi Biotec). Lysates were cleared by two subsequent centrifugation steps at 300 g for 10 min and then 15,000 g for 60 min. 3.6 mg of total protein extracted from tissue and cell-line material was used for subsequent immunopeptidome analysis as determined with Pierce BCA protein assay kit (Thermo Scientific).

### 2.4. MHC-associated peptide purification

One mg per sample of mouse anti-H-2 Kb/Db antibody (ATCC HB-51) was bound and cross-linked to Protein A beads (GE healthcare) using dimethyl pimelimidate (DMP, Sigma). Beads were incubated with cleared lysates overnight, and then washed twice with 150 mM NaCl, once with 450 mM NaCl in 50 mM Tris, pH 8.0, and finally washed with 50 mM Tris, pH 8.0, before elution of peptides and MHC complex components with 10% acetic acid. Peptides were further purified from larger protein components by preparative HPLC on an Ultimate 3000 system (Fisher Scientific). Material was separated on a ProSwift RP-1S 4.6 x 50 mm column (Thermo Scientific) by applying a linear gradient of 2-35% acetonitrile in 0.1 formic acid in water over 10 min. Alternating fractions that did not contain beta-2-microglobulin or any HLA chain were pooled into two final fractions, and further concentrated and kept at −20°C prior to MS analysis [19].

### 2.5. Liquid chromatography tandem mass spectrometry

Mouse MHC samples and all tryptic digested lysates were analysed on a TripleTOF 5600 (SCIEX) coupled to an Eksigent ekspert nanoLC 400 cHiPLC system. Each sample was resuspended in 20 μL 1% acetonitrile, 0.1% formic acid in water. Peptides were separated on an ekspert nanoLC 400 cHiPLC system (Eksigent) supplemented with a 15 cm x 75 μm ChromXP C18-CL, 3 μm particle size using a 60 min linear gradient from 8%-35% acetonitrile in 0.1% formic acid in water at a flow rate of 300 nL/min. Peptides were introduced to a TripleTOF 5600 mass spectrometer, and collision-induced dissociation fragmentation using ramped collision energy was induced on the 30 most abundant ions per full MS scan using unit isolation width 0.7 amu and acquisition time of 120 ms. All fragmented precursor ions were actively excluded from repeated selection for 15 s.

### 2.6. Mass spectrometry data analysis

Analysis of raw data was performed using Peaks 7.5 software (Bioinformatics Solutions) Spider searches. Sequence interpretation of MS/MS spectra was carried out using databases containing all reviewed mouse entries in Uniprot (March 2016, 16,773 entries). Motif analysis of common amino acids in peptide sequences was performed using WebLogo 3.5 (weblogo.threeplusone.com). Peptide clustering was performed with GibbsCluster-2.0 [20], and peptide binding predictions were performed using NetMHC 4.0 [21].

For MHC peptide samples, peptides with a length of less than 7 amino acids were excluded from the analysis results. Spider searches were carried out allowing 2 modifications per peptide. A −10lgP score cutoff of 15 was chosen and false discovery rates (FDR) were estimated using a decoy database search approach. FDR ranged from 2.7 to 6.4% in this analysis.

The mass spectrometry proteomics data have been deposited to the ProteomeXchange Consortium via the PRIDE partner repository with the dataset identifier PXD018459 and 10.6019/PXD018459.

### 2.7. ChAdOx1 and MVA viral vector construction

DNA sequences encoding mouse thymosin beta 4 (Thyβ4) and thymosin beta 10 (Thyβ10) antigens (NCBI Ref Seq: X16053.1 and NM_001190327.1) and fusion constructs of these antigens to the full length mouse invariant chain (CD74) (NCBI RefSeq: NC_000084.6) were obtained from GeneArt (Life technologies, Paisley, UK). The fusion Thyβ4-CD74 and Thyβ10-CD74 constructs were designed as an in-frame fusion of N-terminus of the thymosin cDNA sequences to the C-terminus of the full length murine invariant chain. The construction of ChAdOx1 vector has been described earlier [22]. Briefly, the full length Thyβ4 and Thyβ10 cDNAs or their fusion constructs under a CMV immediate early promoter were sub-cloned from a pENTR plasmid into the E1 locus of the pBAC ChAdOx1-DEST genomic clone by *in vitro* site-specific recombination (GatewayTM cloning). The constructs were then used to transfect HEK293A cells to generate the recombinant adenoviruses expressing these antigens. The MVA-GFP shuttle vector drives the expression of Thyβ4 and Thyβ10 under the p7.5 early/late promoter inserted at the thymidine kinase locus of MVA and the GFP from the fowlpox FP4b late promoter. The plasmids were transfected into MVA-infected primary chick embryo fibroblasts (CEFs; Institute for Animal Health, Compton, UK) and recombinant viruses were isolated by selection of GFP-positive plaques, amplified, purified over sucrose cushions and titred in CEFs according to standard practice. The integrity, identity and purity of the viruses were confirmed by PCR analysis.

### 2.8. Analysis of Thyβ4 and Thyβ10 gene expression

Thyβ4 and Thyβ10 mRNA expression in thymi of C57BL/6 mice and TRAMP-C1 cells was detected by semi-quantitative reverse transcription PCR. Total RNA was extracted from thymic tissue and TRAMP-C1 cells using RNeasy Plus minikit (Qiagen) according to the manufacturers’ instructions. A total of 1-2 μg of RNA was used to synthesize the first single-strand cDNA using QuantiTect Reverse Transcription kit (Qiagen). For RT-PCR amplification, the following priming pairs have been used: Thyβ4 forward primer ACAAACCCGATATGGCTGAG and Thyβ4 reverse primer GCCAGCTTGCTTCTCTTGTT; Thyβ10 forward primer GAAATCGCCAGCTTCGATAA and Thyβ10 reverse primer TTCACTCCTCTTTTCCTGTT; β-actin forward primer GGCATCCTCACCCTGAAGTA and β-actin reverse primer AGCACTGTGTTGGCGTACAG. Transcripts were amplified using 98 °C denaturation, 60 °C annealing and 72 °C extension for a total of 35 cycles.

### 2.9. *In vivo* studies: immunogenicity and tumour models

*In vivo* studies of immunogenicity and efficacy of Thyβ vaccines were carried out as follows. A dose of 10^8^ infectious units (IU) of ChAdOx1 or 10^7^ plaque forming units (PFU) of MVA expressing Thyβ4, Thyβ10 and the respective CD74-fused antigens were diluted in DPBS and injected intramuscularly (50 μl per animal) in C57BL/6 male mice. Alternating immunisations with ChAdOx1 and MVA vectors were performed at 3 week intervals. Alternatively, a protocol of weekly immunisations was performed at a dose of 10^7^ IU of ChAdOx1 and 10^6^ PFU of MVA.

Efficacy of the vaccines was assessed in a transplantable tumour model. For this purpose, C57BL/6 male mice were prime-boosted as described above prior the subcutaneous (s.c.) inoculation of 2×10^6^ TRAMP-C1 cells into the flank in a total volume of 100 μl DPBS. Alternatively, 2×10^6^ TRAMP-C1 cells were injected s.c. and immunisations initiated only at the appearance of palpable tumours. Tumour growth was monitored 3 times weekly and tumour volume was calculated follows: length (mm) x width^2^ (mm) x 0.5. Mice were sacrificed when tumour size reached 10 mm in any direction.

### 2.10. Measurement of thymosin specific immune responses by IFN-γ Elispot

An *ex vivo* IFN-γ Elispot assay was performed using Multiscreen IP ELISPOT plates (Millipore) and mIFN-γ Elispot kit (Mabtech) [23]. Peptide libraries of murine Thyβ4 and Thyβ10 proteins were synthesized by Mimotopes (UK) and consisted of 15-mer peptides overlapping by 10 amino acids spanning the whole protein length. Individual peptide pools or a mix of all pools at a final concentration of 5 μg/ml were used to stimulate PBMCs or splenocytes for 18-22 hours prior to detection of spot-forming cells (SFCs). In some experiments, cells were also stimulated with 5 μg/ml of Thyβ10 polypeptide (Mimotopes). ELISPOT plates were counted using an AID automated ELISPOT counter (AID Diagnostika GmbH) using identical settings for all plates.

### 2.11. Statistical analysis

Each *in vivo* experiment presented in this manuscript is representative of at least 2 replicate experiments using a minimum of 6 animals per group. Unless stated otherwise, median values ± SEM are shown. Area under curve analysis was performed to compare tumour growth kinetics in control versus Thyβ10-vaccinated mice. For comparisons between groups, a one-way ANOVA with post-test comparison was performed. Statistical significance in survival experiments was evaluated by the log-rank test. Differences were considered statistically significant for two-sided *p* values < 0.05. Calculations were performed using GraphPad PRISM package (v6).

## 3. Results

### 3.1. Identification of H-2-associated peptides in primary mouse prostate tumour and the TRAMP-C1 prostate tumour cell line in comparison to benign prostate tissue

For analysis of cancer-specific antigens in mice, we chose the transgenic adenocarcinoma of the mouse prostate (TRAMP) model which closely mimics the development of human prostate cancer [24]. TRAMP male mice uniformly develop prostate tumours due to expression of SV40 Tag oncogene under the control of a prostate-specific androgen-regulated promoter. By 22-25 weeks of age the tumours progress to advanced prostatic adenocarcinoma and mice die of metastatic disease 10-20 weeks later. In our experiment, a malignant prostate gland was excised from a TRAMP male mouse at the age of 24 weeks. For a benign control, we collected and pooled prostate glands from age-matched wild type mice of C57BL/6 background to equal the weight of the tumour sample. In order to compare the repertoire of tumour-specific antigens presented on the cell surface of a primary prostate tumour with a cultured tumour cell line, the murine prostate TRAMP-C1 cell line established from a TRAMP tumour [25] was analysed in parallel. To increase MHC class I expression in this cell line, TRAMP-C1 cells were treated with mouse IFN-γ at 100U/ml for 48 hours before harvesting (Supplementary Figure S1).

C57BL/6 mouse strain has the MHC H-2K^b^, H-2D^b^ haplotype. H-2-associated peptides were purified by immunoprecipitations of the MHC class I-peptide complexes with the HB-51 antibody using equal amounts of total protein extract from the prostate tissue samples and the TRAMP-C1 cell line to allow for quantitative comparison of peptide abundance between the samples. We identified 675 peptide sequences from benign prostate tissue, 1034 peptide sequences from malignant prostate tissue and 1493 peptide sequences from the TRAMP-C1 cell line (Figure 1A). The peptides showed general characteristics of MHC class I-bound peptides: 80, 55 and 77% of peptides had a length of 8 or 9 aa in the respective samples (Figure 1B), and most of the peptides exhibited a charge state of 2 (Figure 1C). Between 72-82% of 8-mer peptides were predicted to bind to H2-K^b^, while 71-79% of 9-mer sequences were predicted to bind to H-2D^b^ across all samples with a NetMHC binding rank score ≤2 (Figure 1D). Both alleles generally show strong peptide length preference [26], which we could also observe in our experiment: 8-mer peptides exhibited the characteristics of H-2K^b^-while 9-mer peptides had motifs specific to H-2D^b^-binding without any prior clustering of peptide sequences (Figure 1E). Gibbs clustering of all 8- and 9-mer peptide sequences, respectively, resulted in clear motifs for either allele in comparison to the known sequence motif as constructed with all peptides deposited in IEDB (Figure 1F). Overall, we identified 687 peptides in tumour tissue only, and 1117 peptides unique to the TRAMP-C1 cell line in comparison to healthy prostate tissue (Figure 1G).

**Figure 1.**
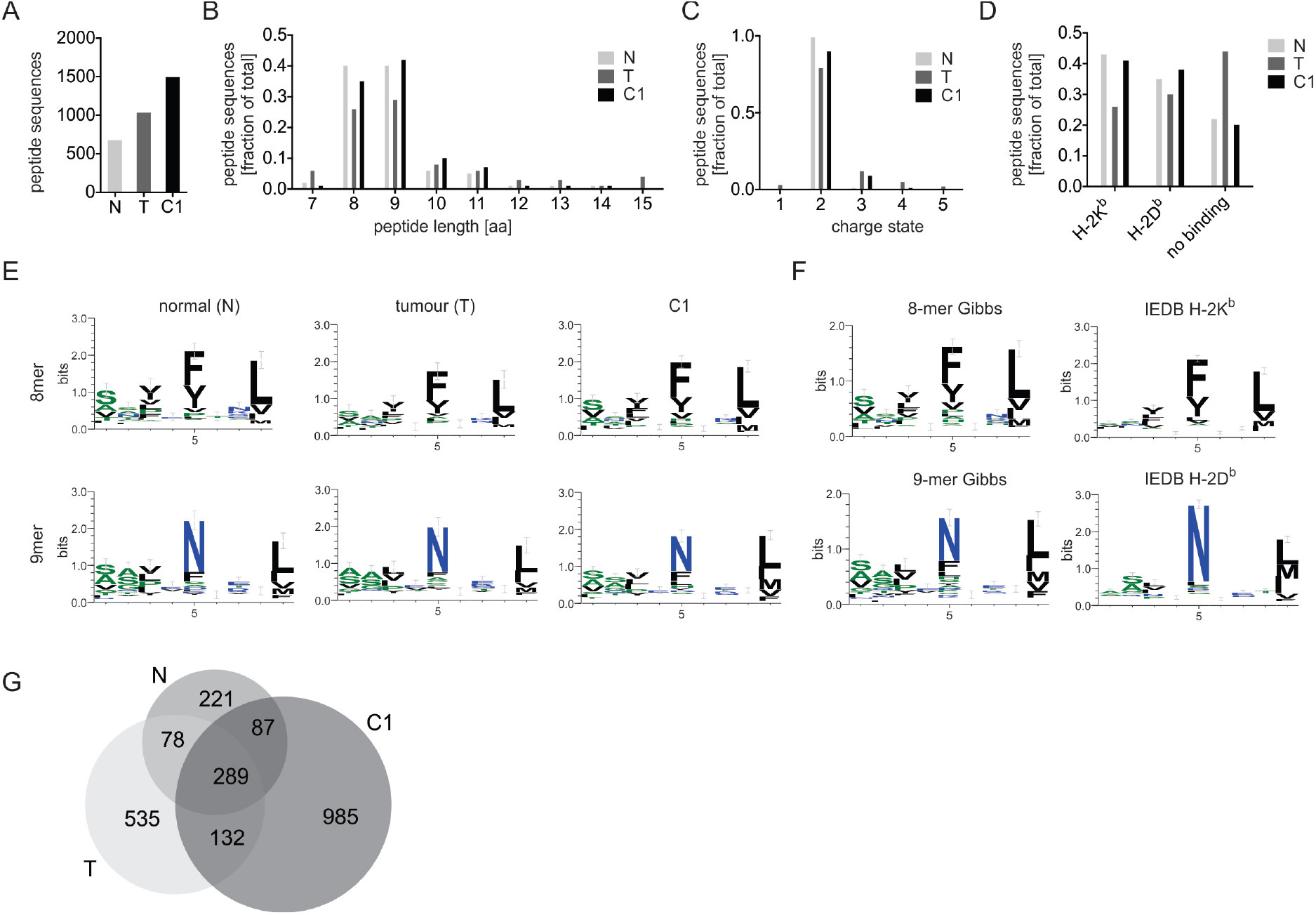
Characteristics of identified peptides from benign mouse prostate, malignant mouse prostate and the TRAMP-C1 cell line. For each sample, the total number **(A)**, length distribution **(B)**, and charge state distribution **(C)** of identified peptide sequences are shown. **(D)** Number of peptides predicted to bind to H-2-Kb and Db using NetMHC 4.0. **(E)** Motifs of the frequency of amino acids are depicted for each sample for 8- and 9-mer peptide sequences. **(F)** Results of Gibbs clustering (left panels) in comparison to motifs from all 8-mer sequences for H-2-Db and all 9-mer sequences for H-2-Kb deposited in IEDB. **(G)** Overlap of peptide sequences between all samples. N= benign mouse prostate; T= malignant mouse prostate; C1= TRAMP-C1 cell line.

### 3.2. Immunopeptidomic analysis reveals presentation of thymosin beta family polypeptides with highest tumour-specific presentation coverages

Thymosin beta family polypeptide Thyβ4 and Thyβ10 were prostate tumour-specific antigens identified with the highest coverage. Thyβ10 was present in both the prostate tumour resection specimen and TRAMP-C1 cells, while Thyβ4 was detected in tumorous prostate tissue only. Both antigens were not detected in normal prostate tissue. In the prostate tumour resection sample, 30 peptides of Thyβ4 (98% protein sequence coverage), and 7 peptides of Thyβ10 (86% protein sequence coverage) were identified. Three peptides of Thyβ10 were identified in the TRAMP-C1 cell line, resulting in 39% protein coverage. The eluted peptides had a length of between 12-26 aa (Figure 2).

**Figure 2.**
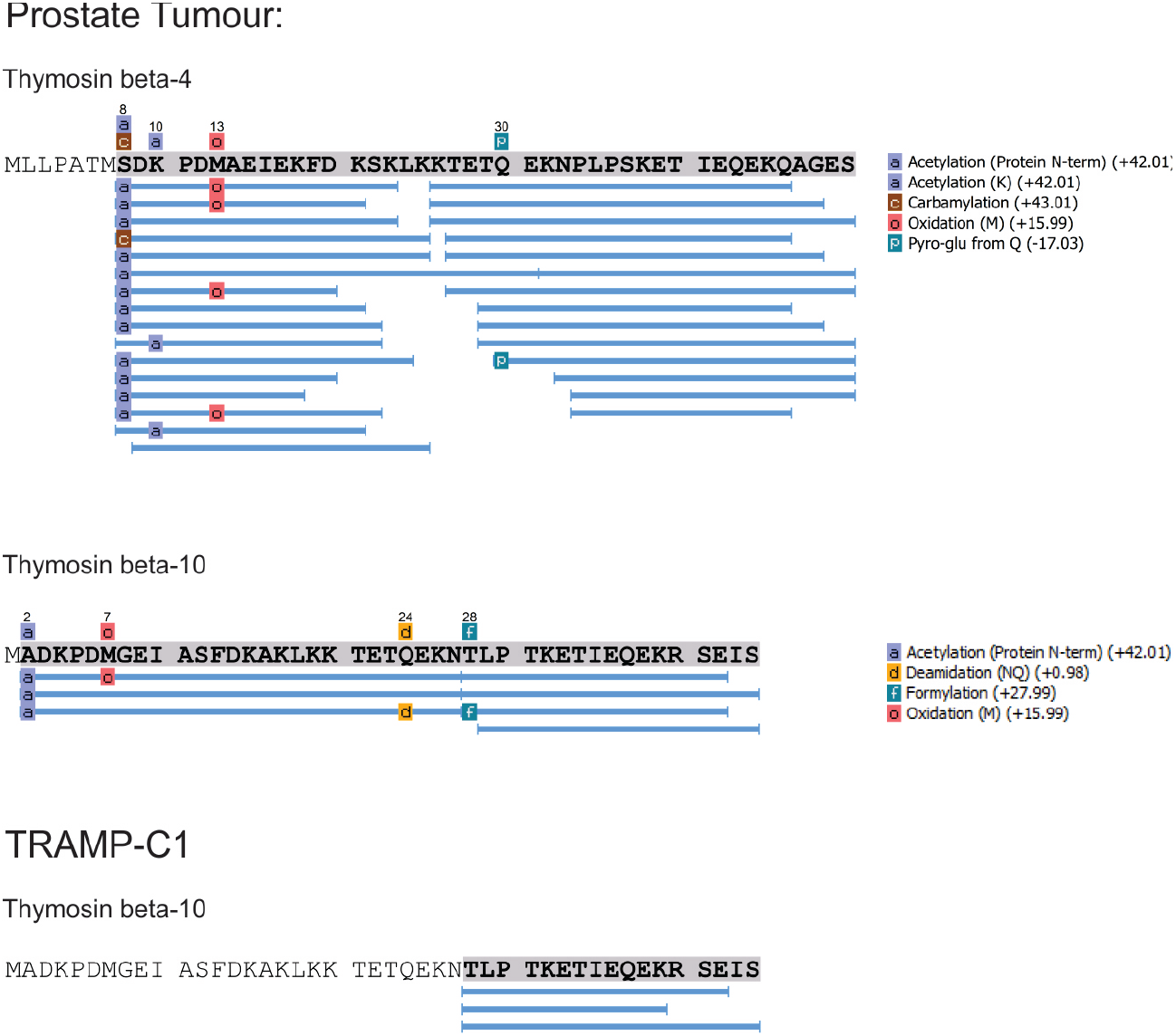
Peptide coverages of thymosin beta family antigens in prostate tumour tissue and the TRAMP-C1 cell line. Each blue line represents an identified peptide sequence. Peptide modifications are indicated as colored squares at the respective residue in the peptide string and modifications are listed in the legend.

### 3.3. Immunogenicity of viral vectors encoding Thyβ4 and Thyβ10 in C57BL/6 mouse strain and their protective efficacy in the TRAMP-C1 transplantable tumour model

Thyβ4 and Thyβ10 polypeptides belong to the family of highly conserved 5 kDa proteins originally isolated from the calf thymus and thought to be a thymic hormone [27]. They are involved in many cellular processes such as cellular motility, angiogenesis, inflammation, cell survival and wound healing. Recently, a role for β-thymosins has been proposed in the process of carcinogenesis as both peptides were detected in several types of cancer and played a key role in facilitating tumour metastasis and angiogenesis [28–32].

These findings along with their exclusive presence on the surface of the TRAMP-C1 tumour detected by MS, prompted us to investigate whether these self-antigens can serve as antigenic targets for a novel prostate cancer vaccine. To this end, firstly we performed *in vivo* experiments to evaluate the immunogenicity of Thyβ4 and Thyβ10 antigens in C57BL/6 mice using ChAdOx1 and MVA recombinant viral vectors. This vaccination platform, developed at our laboratories over a decade ago, has been shown to be the most powerful approach for inducing polyfunctional protective T cell responses against antigens from diverse human pathogens in clinical trials [33–38], and more recently it has been shown to break tolerance to a tumour-associated self-antigen in preclinical studies [39, 40]. In addition, we have also generated ChAdOx1 and MVA vectors encoding Thyβ polypeptides linked to the mouse MHC class II-associated invariant chain (CD74), as such fusion constructs have been shown to be more immunogenic compared to native antigens [41–45]. To assess immunogenicity of the native antigens and Thyβ-CD74 fusion constructs, C57BL/6 mice were prime-boosted at 3 week interval with 10^8^ IU of ChAdOx1 and 10^7^ pfu of MVA expressing the respective antigens. After each vaccination, an induction of Thyβ-specific T cell responses was assessed by an *ex vivo* IFN-γ ELISPOT assay using a pool of Thyβ peptides covering the entire polypeptide length for PBMC stimulation. As a result, none of the constructs was able to break T cell tolerance to Thyβ self-antigens. Thyβ-specific T cells have not been detected by IFN-γ ELISPOT after short-term (12 days) *in vitro* culture of splenocytes from vaccinated mice in the presence of Thyβ peptide antigens either (data not shown).

The complete absence of antigen specific T cell responses was not surprising given their high expression level in the murine thymus. Although breaches in central tolerance can occur [46], both Thyβ proteins were found to be expressed in thymic tissue at the mRNA level (Figure 3A), suggesting an established central tolerance to thymosin antigens in C57BL/6 mice. The presence of Thyβ4 and Thyβ10 mRNA was also confirmed in the TRAMP-C1 tumour cell line, in the TRAMP prostate and in prostate tissue isolated from age-matched C57BL/6 mice (Figure 3A).

**Figure 3.**
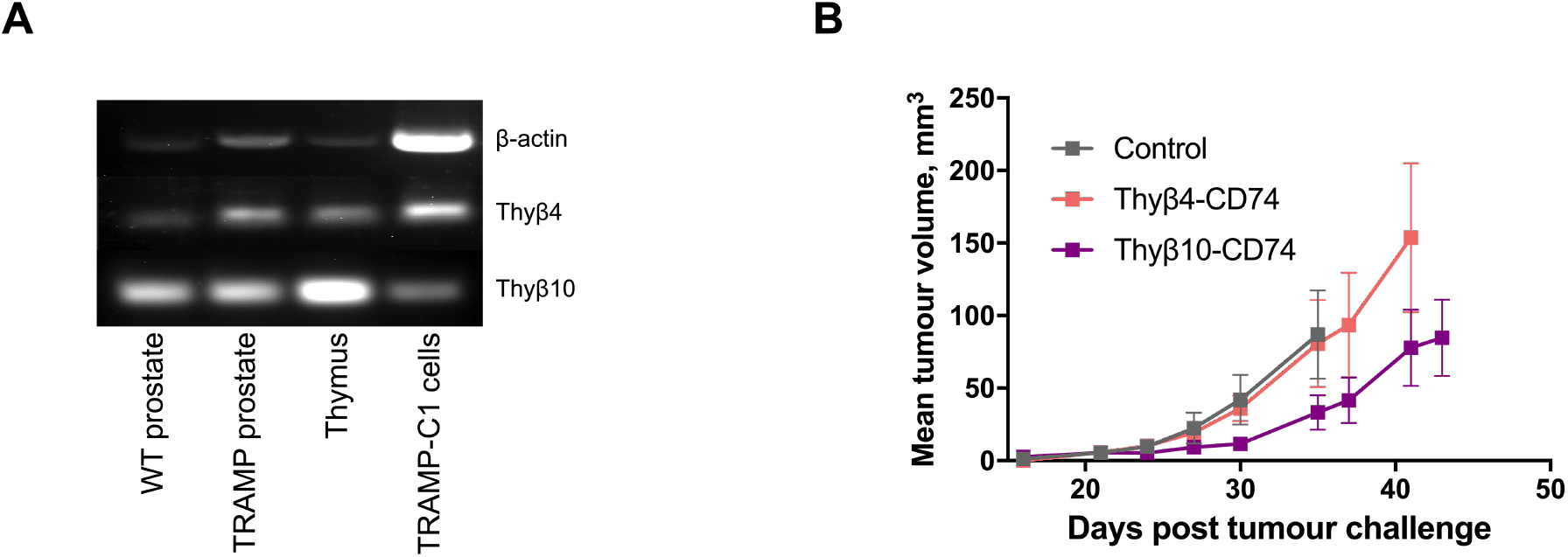
Thymosin beta family protein expression in murine tissue and their efficacy as prophylactic vaccines in a transplantable tumour model. **(A)** Comparative expression of Thyβ4 and Thyβ10 mRNA transcripts in age-matched wild type [47] and TRAMP prostates, murine thymus and TRAMP-C1 cells. cDNA was amplified with primers specific for β-actin, Thyβ4 and Thyβ10 as indicated in Materials and Methods. **(B)** C57BL/6 mice were immunised i.m. at 3 week intervals with 10^8^ IU of ChAdOx1 and 10^7^ PFU of MVA expressing Thyβ4-CD74, Thyβ10-CD74 or equivalent doses of GFP expressing vectors (control) and subsequently challenged with TRAMP-C1 cells subcutaneously. Tumour size was measured 3 times per week and volumes were calculated as described in Materials and Methods. Mean tumour volumes ± SEM comparison between the 3 groups. Representative data of 3 biological replicate experiments are shown. Bars represent median.

In order to understand whether vaccination against Thyβ4 and Thyβ10 could mediate tumour protection through mechanisms other than antigen-specific T cells, we employed tumour challenge studies using the TRAMP-C1 cell line. For this purpose, in a preventive vaccination setting, mice were immunised with 10^7^ IU of ChAdOx1 and 10^6^ pfu of MVA vectors encoding Thyβ-CD74 fusion constructs one week apart and were inoculated subcutaneously with 2×10^6^ TRAMP-C1 cells one day post MVA boost. Control mice were left unvaccinated before tumour challenge. Tumour volumes were monitored and measured at regular intervals throughout the experiment. As shown in Figure 3B, the tumour growth rate in Thyβ4-CD74 vaccinated mice was comparable to the unvaccinated group, while we could observe a slower TRAMP-C1 tumour progression in Thyβ10-CD74 vaccinated mice. Based on these encouraging results, we then set out to perform therapeutic efficacy experiments using exclusively Thyβ10 vaccines. Mice were challenged subcutaneously with 2×10^6^ TRAMP-C1 cells and, upon establishment of palpable tumours, were randomised in three groups. Two groups of mice started a weekly immunisation schedule with 10^7^ IU of ChAdOx1 and 10^6^ pfu of MVA expressing either Thyβ10 or Thyβ10-CD74. Control mice were prime-boosted accordingly using viral vectors expressing an irrelevant antigen. Figure 4A shows individual mouse tumour growth curves in the 3 experimental groups and Figure 4B shows mean tumour volumes in these 3 groups of mice. Only a trend for reduced tumour volume and improved survival was found in the Thyβ10 vaccinated group as compared with control mice (Figure 4C, D). Therapy of established tumours with vaccines expressing Thyβ10-CD74 resulted in a significant reduction of tumour outgrowth in comparison to control mice, and also in a significant survival benefit (Figure 4C, D).

**Figure 4.**
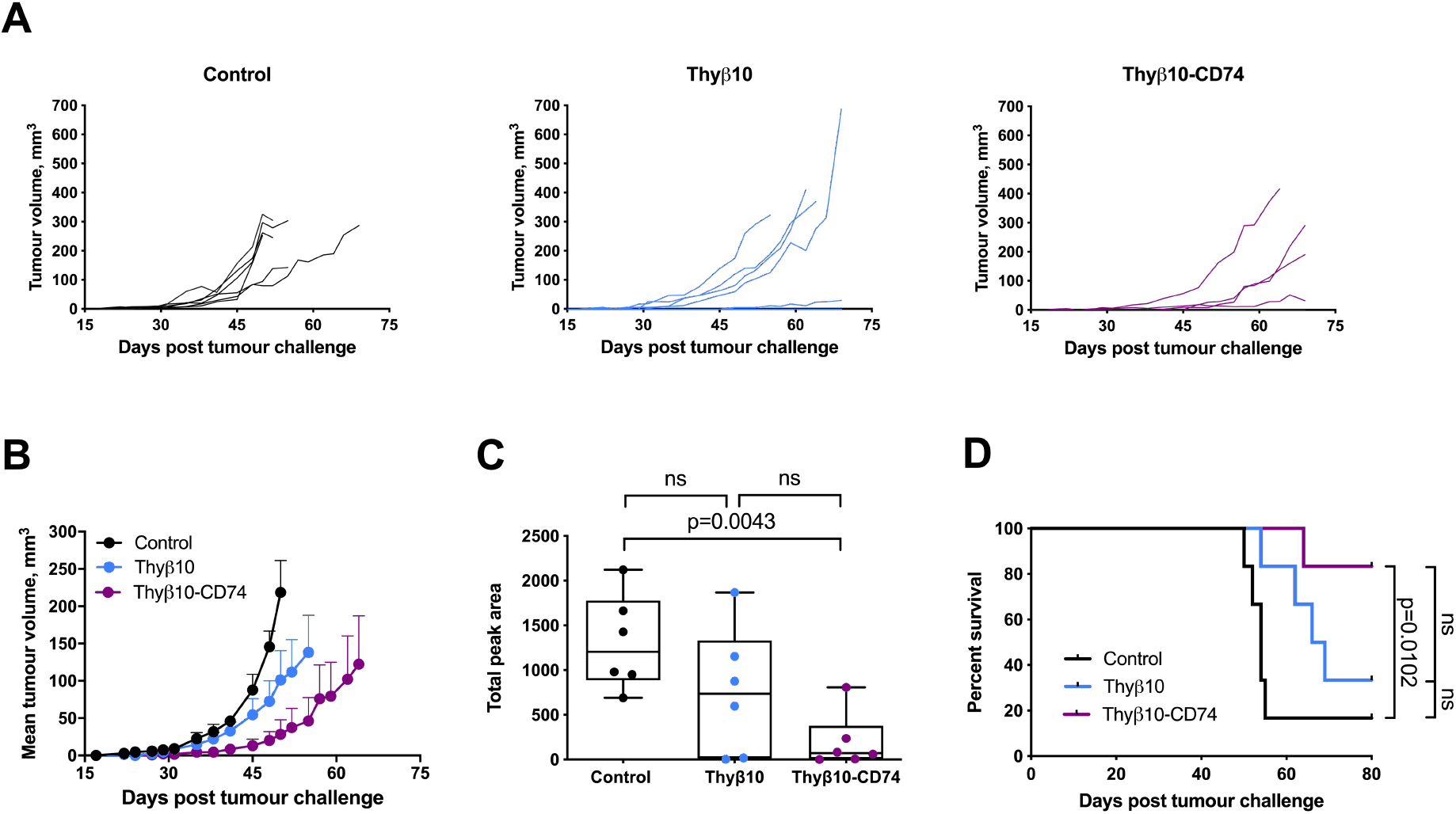
Therapeutic Thyβ10-CD74 vaccination reduces tumour growth rate and improves survival in a transplantable tumour model. 6-week old male C57BL/6 mice were challenged subcutaneously with TRAMP-C1 cells and randomised in 3 groups. At the appearance of palpable tumours, groups were immunised intramuscularly at 1 week intervals with alternating doses of 10^7^ IU of ChAdOx1.Thyβ10 and 10^6^ PFU of MVA.Thyβ10, or with 10^7^ IU ChAdOx1.Thyβ10-CD74 and 10^6^ PFU of MVA. Thyβ10-CD74. Control group was immunised weekly with equivalent doses of vectors expressing an irrelevant antigen. Tumour size was measured 3 times per week and volumes were calculated as described in Materials and Methods. Representative data of 2 biological replicate experiments are shown. **(A)** Individual mouse tumour growth curves in the 3 experimental groups. **(B)** Mean tumour volumes ± SEM comparison between the three groups. **(C)** Assessment of vaccine efficacy by area under the curve (AUC) analysis at day 50 post tumour challenge. Bars represent median. P values are shown. **(D)** Kaplan Meyer survival curves of the three groups of vaccinated mice. Logrank (Mantel-Cox) test P value is shown. P values are shown. ns=not significant.

To investigate further the possible mechanisms of protection by the Thyβ10-CD74 vaccine, PBMCs and splenocytes of vaccinated mice were subjected to flow cytometric analysis to measure secretion of effector cytokines other than IFN-γ. As a result, no significant increase in the production of IFN-γ, IL-2, TNF-α or CD107a was detected in CD4+ and CD8+ T cells. The protective role of the antibody response has also been assessed through measurement of anti-thymosin antibody titres of pooled sera of vaccinated mice compared to control mice in an ELISA assay. No Thyβ10-specific antibody responses were detected in Thyβ10 or Thyβ10-CD74 vaccinated mice either (data not shown).

## 4. Discussion

Although the cancer immunotherapy field has been recently reinvigorated by the emergence of checkpoint inhibitors, the most clinically advanced antigen-specific cancer vaccines have had only limited clinical success [1, 4]. One of the many reasons for such inadequate outcome for cancer vaccines is the challenge of identifying ideal vaccine target antigens. Immunopeptidome analysis of tumour cells by mass spectrometry is a new and promising strategy for discovery of tumour-specific antigens and, therefore, for the development of cancer vaccines, as it offers the advantage of identifying tumour-specific proteins that are naturally processed and presented on the surface of the tumour cells in the context of MHC molecules.

In the current study, we have performed a comparative analysis of the natural peptidome in benign and malignant murine prostate gland. We have focused on the comparison of MHC class I-associated peptides presented in the TRAMP tumour sample with peptides presented in healthy mouse prostate tissue, with the aim of identifying proteins that are upregulated or presented only in the cancerous tissue. We have also included in the analysis TRAMP-C1 cells cultured in the presence of IFN-γ in order to upregulate MHC class I expression, thereby increasing the yield of eluted peptides [48], since these cells are known to have generally very low MHC class I expression [24]. Our comparative analysis of the immunopeptidome in normal prostate tissue and TRAMP tumour led to the identification of 2327 peptide sequences from MHC enriched samples, of which 1652 peptides were only detected in either malignant prostate gland or the TRAMP-C1 cell line. Thyβ family antigens 4 and 10 had the highest coverage in prostate tumour samples, while they were not detected in healthy tissue. Identified peptides ranged between 11-26 aa, which is unusually long for presentation by class I molecules which is normally restricted to 8-12mer sequences. Measured sequences were not predicted to bind to the MHC I alleles H-2D^b^ and H-2K^b^ as determined by NetMHC searches (data not shown), however longer presented peptides have been previously observed in the context of human class I presentation following IFN-γ treatment [49]. Another possibility could be that thymosins localise to the membrane and/or specifically interact with MHC molecules in tumour tissue. Finally, we cannot exclude the possibility that these peptides are not directly presented by MHC molecules, but that these antigens co-precipitated in the IP experiment. We have previously observed co-precipitation of membrane-bound proteins, in particular MHC class II proteins and subsequent co-purification of MHC class II antigenic ligands in the context of MHC class I IP [50]. Also, we have observed an upregulation of the MHC class II expression on the TRAMP-C1 cells following IFN-γ treatment (Supplementary Figure S1). Therefore, the longer peptides could potentially be originating from MHC II molecules and future studies should aim to understand the detailed presentation and recognition requirements for the detected antigens in the TRAMP autochthonous tumour model.

Importantly, Thyβ4 and Thyβ10 polypeptides are 100% homologous in humans and mice, highlighting a potential opportunity for clinical relevance and translation. Thymosin family members have been long known to be involved in physiological processes such as angiogenesis, cell motility and wound healing [51, 52]. Thymosin family members are also involved in tumour invasion and cell migration; in addition, expression of members of this family correlates with tumour progression and poor prognosis in different tumour types [29–32, 53, 54]. Although Thyβ4a and Thyβ10 antigens are ubiquitously expressed in human tissues on the RNA level (https://www.proteinatlas.org), the presence of thymosin transcripts in many tissue types does not necessarily correlate with the expression of these polypeptides at the protein level, which alleviates concerns about a potential vaccine-induced off-target toxicity. As an example, the tumour-associated antigen 5T4, that has been extensively studied in preclinical and clinical settings as a cancer vaccine target, is also ubiquitously expressed in human and mouse tissues on the RNA level; however, we have never observed any adverse events related to autoimmunity or collateral damage to normal tissue following vaccine-induced 5T4-specific T cell immune responses in mice or humans [40, 55]. We have therefore selected Thyβ4 and Thyβ10 as target antigens for a cancer vaccine in the C57BL/6 mouse strain. At first, we have engineered the ChAdOx1 and MVA viral vectors to express Thyβ4 and Thyβ10. The combination of these viral vectors in a heterologous prime-boost regimen is a potent inducer of CD8+ T cell responses, both in infectious disease and in cancer settings. The two antigens were also linked to CD74, since tethering of the invariant chain to native antigens has been long known to accelerate, enhance, broaden and prolong CD8+ T cell responses and, importantly, to markedly delay tumour growth compared to unmodified antigens in mouse models [41, 42]. Despite deployment of potent inducers of CD8+ T cell response, such as ChAd-MVA regimen and CD74 fusion, we could not observe antigen-specific IFN-γ responses *ex vivo* and even after a short-term *in vitro* culture with Thyβ polypeptides and low dose recombinant mIL-2. In line with the lack of immunogenicity of the identified thymosin proteins, we have found high expression of thymosin at the mRNA level in the murine thymus. This finding supports our hypothesis that thymosin-specific T cell receptor repertoire was effectively eliminated during negative selection in the thymus. Of note, the vaccines have not induced a thymosin-specific antibody response in our experiments either.

Nevertheless, we set out experiments to investigate the tumour-protective efficacy of the vaccines in the TRAMP-C1 transplantable tumour model. In a prophylactic setting, we observed only a modest effect of the Thyβ10-CD74 targeted vaccine on tumour growth. Surprisingly, even in the absence of IFN-γ responses in the periphery and modest prophylactic efficacy, Thyβ10-CD74 vaccination translated in a significant delay in tumour growth and highly significant improvement of survival in mice when administered in a therapeutic setting. Potential mechanisms underlying tumour protection in our experiments remain to be elucidated. As an example, neoantigen-based cancer vaccines can elicit tumour rejection in a CD8-dependent manner, however, they may still not be able to elicit CD8+ T cell responses measurable *in vitro* [56]. Other studies, rather than focusing on cytotoxic activity and IFN-γ secretion to demonstrate cancer vaccine anti-tumour effect, have concentrated on the proliferation capacity and expression of activation markers on T cells. Also, alterations of T cell populations, such as regulatory T cells, in the periphery and within the tumour microenvironment have been examined [57, 58]. We believe the same analysis could help us define in more details the anti-tumour efficacy of Thyβ10 vaccination in future, although our preliminary flow cytometry data have not highlighted any particular marker. Open questions include the identification of immune cell subsets induced by vaccination that are directly involved in tumour growth delay, their functionality and cytotoxic mechanisms. There is some evidence indicating that components of the immune system other than CD8+ T cells can play an important role in tumour clearance [59]. Likewise, we can speculate that in our model anti-tumour mechanisms could involve Fas/Fas-ligand engagement, TNF-related apoptosis or antibody-dependent cell-mediated cytotoxicity, however, the presence of the invariant chain seems to be critical.

Recently, a novel population of memory CD8+ T cells, the tissue resident memory T cells (T) has emerged as a key player in the efficacy of cancer vaccines [60]. As these cells are committed to reside permanently in tissues without recirculating in blood, it may explain why we could not detect peripheric Thyβ-specific T cells in tumour-bearing mice. In line with this notion, Sipuleucel-T led to local tumour infiltration of CD8+ T cells without the detection of antigen specific CD8+ T cells in the blood [61]. On the contrary, most cancer vaccines failed to demonstrate clinical benefit even in the presence of anti tumour CD8+ T cells in the blood [62–64]. Trm cells are found in diverse human solid cancers where their presence correlate with improved prognosis [65, 66] and they can protect against tumour challenge in mice [67]. The mechanisms through which these cells mediate cancer protection still remain poorly understood. It has been demonstrated that mucosal immunisations were more efficient than the conventional systemic routes (intramuscular, subcutaneous, etc.) to elicit Trm cells at the mucosal tumour site [67]. However, the development of Trm cells may be initiated in the lymph nodes by a specific type of dendritic cells [68], and cross-priming of dendritic cells seems to be critical for the generation of resident memory in the tissues [69]. Local inflammatory signals after a systemic vaccination (prime-pull protocol) could also favour the induction of Trm cells [70, 71]. In light of the above and given that the fusion of Thyβ10 to CD74 was crucial for the tumour growth delay, it is plausible that such fusion promoted the induction of tumour-protective Trm cells in our experiments.

## 5. Conclusions

In summary, the herein presented results represent one of the very few successful examples of tumour target discovery in preclinical immunotherapy settings, from the identification to the *in vivo* assessment of the candidate antigen. In the expanding field of personalised medicine, peptidome analysis of tumours and identification and exploitation of tumour-specific antigens have high potential to improve the efficacy of immunotherapy, especially if MS analysis is combined with next generation sequencing for the identification of tumour neoantigens [72]. We are currently applying the same analysis to human prostate tissue samples with the aim of clinical translation, with several considerations to be raised. One of them is that LC-MS approach does not allow to ensure that the identified peptides originate from tumour cells only, as biopsies, even when tumour targeted, can be composed of a mixture of populations, including normal cells [73]. It is important to highlight that a MS-based approach of this kind in humans will also face a broader heterogeneity between patients, not only influenced by the nature of the tumour and tumour mutational burden, but also by the HLA type and the natural status and efficacy of the immune system.

## Supplementary Materials

Supplementary Materials and Methods: MHC I and MHC II expression on TRAMP-C1 cells by flow cytometry; In vitro culture of splenocytes; Immune cell phenotyping by flow cytometry; ELISA. Supplementary Figure S1. MHC I and MHC II upregulation on TRAMP-C1 cells upon IFN-γ treatment

## Author Contributions

FC, NT, IR designed and performed research, analysed data, and wrote the manuscript. FC, JM, and EP contributed in conducting experiments. AVSH provided scientific input and assisted with the preparation of the manuscript.

## Funding

This work was supported by the European Union’s Seventh Framework Programme under grant agreement No. 602705.

## Acknowledgements

Raw MS data were acquired in the Target Discovery Mass Spectrometry Laboratory run by Benedikt Kessler.

## Competing Interests

The authors declare that the research was conducted in the absence of any commercial or financial relationships that could be construed as a potential conflict of interest.

## Data Availability Statement

All datasets generated for this study are included on the Article/Supplementary Materials. MS data were submitted to the public online data repository PRIDE.

## Notes

### Competing Interest Statement

The authors have declared no competing interest.

