## Supplementary for "The viral vectored vaccine targeting Thymosin β10 controls tumour growth in a murine model of prostate cancer": Supplementary Materials and Methods.docx

**MHC I and MHC II expression on TRAMP-C1 cells by flow cytometry**

TRAMP-C1 cells cultured in the presence or absence of recombinant mIFN-γ (100U/ml for 48 hours) were analysed for MHC I and MHC II expression by flow cytometry. Briefly, cells were harvested and, after Fc block, incubated with the purified anti-H-2 Kb/Db antibody (purified from HB-51 hybridoma supernatant) followed by FITC conjugated anti-mouse IgG antibodies, or incubated with FITC anti-mouse I-A/I-E (Biolegend). Dead cells were discriminated by Live/Dead Near IR fixable staining (Life technologies). All sample measurements were performed on a BD LSRII^TM^ analyzer and data analysed with FlowJo software (Treestar).

***In vitro* culture of splenocytes**

Splenocytes isolated from Thyβ and Thyβ-CD74 vaccinated mice and from naïve control mice were cultured *in vitro* to expand Thyβ4 or Thyβ10 specific T cells. Briefly, 3 x 10^6^ freshly isolated splenocytes/ml were plated in complete medium with or without total Thyβ4 or Thyβ10 peptide pool (5μg /ml). Every 3-4 days the medium was replaced and recombinant mouse IL-2 (100U/ml) added to the cultures. Splenocytes were harvested after 2 weeks and incubated overnight at 37 °C before performing a standard IFN-γ Elispot assay.

**Immune cell phenotyping by flow cytometry**

Mouse PBMCs or splenocytes were stimulated *ex vivo* with 7.5 µg/ml of pooled Thyβ10 peptides or with Thyβ10 polypeptide synthesized by Mimotopes (UK) in the presence of CD107a PE-Cy7 (clone 1D4B) for 6 hours. One µg/ml each of Golgi-Plug and Golgi-Stop (BD) was added in the last 4 hours of stimulation. Following stimulation, cells were incubated with unconjugated anti-CD16/32 to prevent non-specific Fc binding. Cells were then labelled with anti-mouse CD4 AlexaFluor700 (clone RM4-5), CD8 PerCPCy5.5 (clone 53-6.7) and Live/Dead Near IR. Cells were then fixed-permeabilized in CytofixCytoperm buffer (BD) and incubated with IFN-γ PE (clone XMG1.2), IL-2 APC (clone JES6-5H4) and TNF-α FITC (clone MP6-XT22) antibodies.

All samples were acquired on a BD LSRII^TM^ analyzer and data analysed with FlowJo software (Treestar).

**ELISA**

Pooled sera of naïve mice or mice vaccinated with viral vectors expressing murine Thyβ10 and the CD74-fused antigen were tested for anti-Thyβ10 antibody responses by enzyme-linked immunosorbent assay (ELISA) using synthetic Thyβ10 polypeptide (Mimotopes, UK) at 2.5ug/ml as a coating antigen. Rabbit polyclonal antibodies to Thyβ10 (Abcam) were used as positive control. Briefly, Thyβ10-coated plates were blocked with DPBS, 2% FBS before incubating with sera dilutions. Anti-mouse or anti-rabbit IgG HRP-conjugated antibodies (Jackson ImmunoResearch Labs) and OPD peroxidase substrate (Sigma) were used to detect and develop Thyβ10-specific antibodies. Endpoint titres in vaccinated mice were calculated as the highest serum dilution that yielded at least a 2 fold greater OD compared to the naïve serum.

**Supplementary Figures**

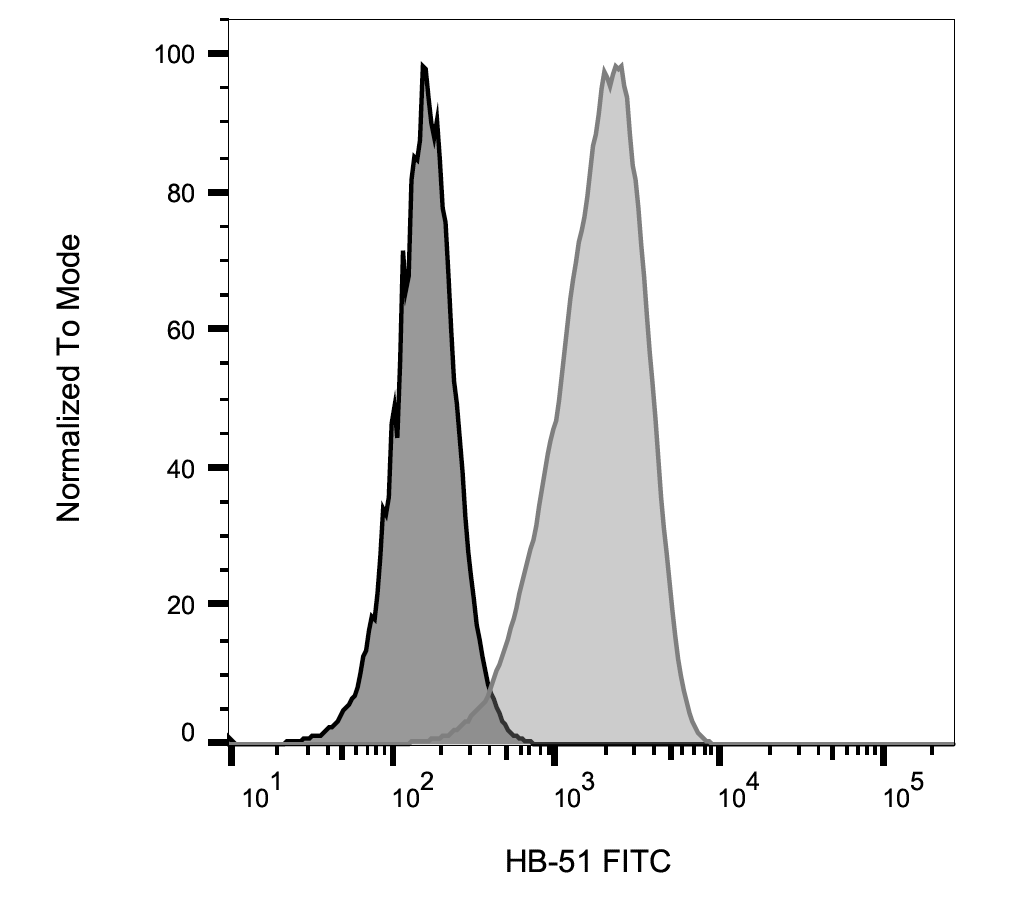

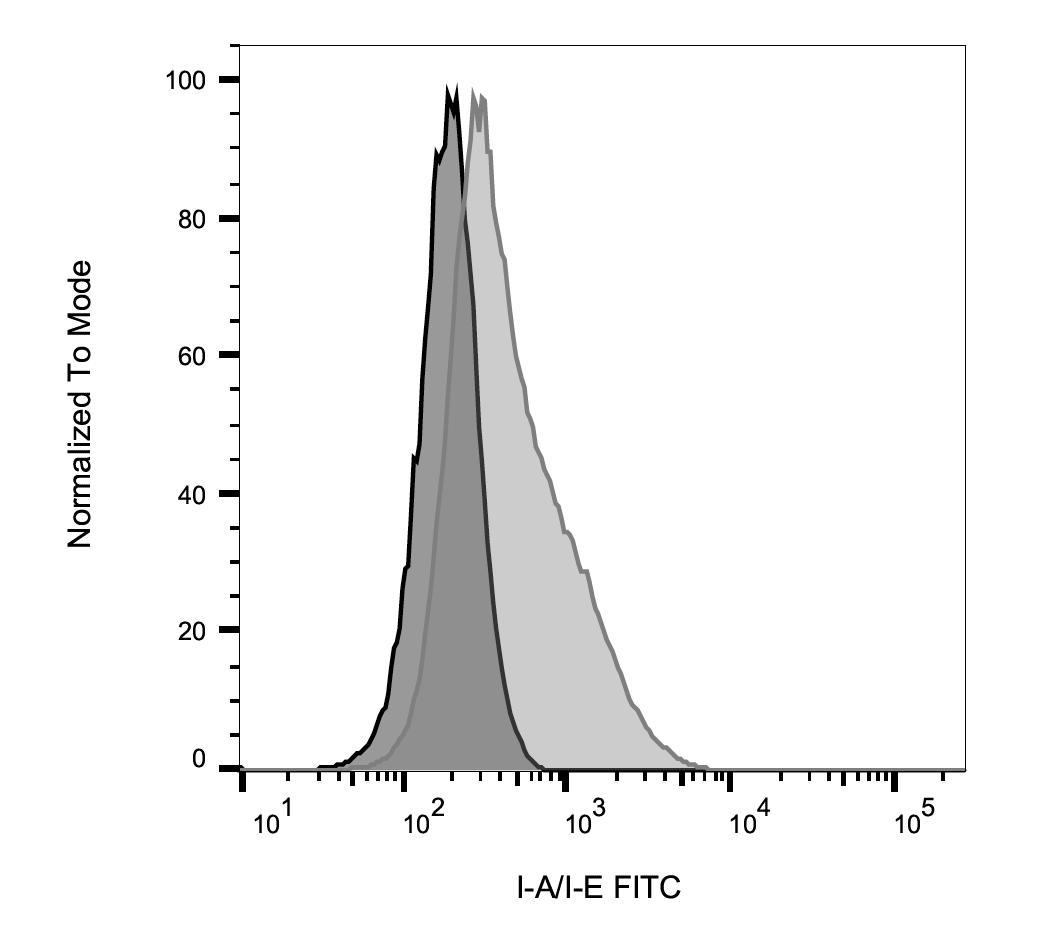

TRAMP-C1

TRAMP-C1+IFN-γ

**Supplementary Figure. S1.** MHC I and MHC II upregulation on TRAMP-C1 cells upon IFN-treatment. TRAMP-C1 cells were cultured for 48 hours in complete medium or in complete medium supplemented with 100 U/mL of recombinant mouse IFN-γ. Cells were subsequently dissociated and incubated with anti MHC I purified from HB-51 hybridoma cells (left panel), or anti MHC II (I-A/I-E) antibodies (right panel). Black histograms represent TRAMP-C1 cells and grey histograms represent IFN-γ treated TRAMP-C1 cells.
